# A core of functional complementary bacteria infects oysters in Pacific Oyster Mortality Syndrome

**DOI:** 10.1101/2020.11.16.384644

**Authors:** Camille Clerissi, Xing Luo, Aude Lucasson, Shogofa Mortaza, Julien de Lorgeril, Eve Toulza, Bruno Petton, Jean-Michel Escoubas, Lionel Dégremont, Yannick Gueguen, Delphine Destoumieux-Garzόn, Annick Jacq, Guillaume Mitta

## Abstract

**Background:** The Pacific oyster *Crassostrea gigas* is one of the main cultivated invertebrate species worldwide. Since 2008, oyster juveniles have been confronted with a lethal syndrome known as the Pacific Oyster Mortality Syndrome (POMS). POMS is a polymicrobial disease initiated by a primary infection with the *herpesvirus* OsHV-1 μVar that creates an oyster immunocompromised state and evolves towards a secondary fatal bacteremia. In the present article, we describe the implementation of an unprecedented combination of metabarcoding and metatranscriptomic approaches to show that the sequence of events in POMS pathogenesis is conserved across infectious environments. We also identified a core bacterial consortium which, together with OsHV-1 μVar, forms the POMS pathobiota. This bacterial consortium is characterized by high transcriptional activities and complementary metabolic functions to exploit host’s resources. A significant metabolic specificity was highlighted at the bacterial genus level, suggesting low competition for nutrients between members of the core bacteria. Lack of metabolic competition might favor complementary colonization of host tissues and contribute to the conservation of the POMS pathobiota across distinct infectious environments.

## INTRODUCTION

Introduced from Asia to a broad range of countries, *Crassostrea gigas* has become one of the world’s main cultivated species. Since 2008, juvenile stages of *C. gigas* have suffered massive mortality events, especially in France [1]. In subsequent years, this so-called Pacific Oyster Mortality Syndrome (POMS) has become panzootic [2]. POMS has been observed in all coastal regions of France [3–5] and numerous other countries worldwide [1,6–10]. Multiple factors contribute to the disease and its severity including seawater temperature, oyster genetics, oyster age, microbiota and infectious agents [11–18]. Thus, dramatic POMS mortality events have coincided with the recurrent detection of *Ostreid herpesvirus* variants in moribund oysters [3–5] as well as bacterial strains of the genus *Vibrio* [19,20].

Recently, integrative molecular approaches have revealed the complex etiology of POMS, which involves an interaction between the viral and bacterial agents involved in the pathosystem [21,22]. Infection by *Ostreid herpesvirus* type 1 μVar (OsHV-1 μVar) is the initial step that leads to an immunocompromised state of oysters. The resulting dysbiosis and bacteremia ultimately result in oyster death [21]. Several bacterial genera are involved in this secondary infection [21]; among them *Vibrio* behave as opportunistic pathogens that cause hemocyte lysis [22]. *Vibrio* species are not the only bacteria that colonize oyster tissues during the secondary bacterial infection. Several bacterial genera, including *Arcobacter, Marinobacterium, Marinomonas*, and *Psychrobium* were also found to massively colonize OsHV-1-infected oysters [21].

The polymicrobial nature of POMS was characterized based on observations in a French Brittany infectious environment [21]. We still ignore whether POMS pathogenesis is conserved in terms of sequence of events and bacterial partners in other regions affected by the disease. In addition, the mechanisms underlying the colonizing capacity of the bacterial consortium have not been elucidated.

In the present study, we investigated whether POMS pathogenesis is conserved across environments, and which biological functions are expressed by the bacterial consortium that causes oyster death. We compared pathogenesis using oyster biparental families that display contrasting phenotypes (resistant or susceptible to POMS) confronted to two infectious environments (the Atlantic Bay of Brest, and the Mediterranean Thau Lagoon). We found that the sequence of events is conserved within both infectious environments: it starts with an intense viral replication in susceptible oysters, followed by a secondary bacteremia caused by a conserved bacterial consortium that results in oyster death. Using metabarcoding and metatranscriptomics, we identified in the present work members of the core pathobiota and characterized their functional response to host environment. We found that each bacterial genus has a reproducible transcriptional response across infectious environments. In particular, translation and central metabolism were highly induced. Results indicate that metabolism might play an important role in tissue colonization, and that metabolic complementarity between members of the core consortium possibly explains the conservation of this assemblage across environments.

## METHODS

### Production of biparental oyster families

*C. gigas* families were produced as described in [21,23]. Briefly, oysters were produced at the Ifremer hatchery in Argenton in March 2015. Three susceptible families (F_11_, F_14_, and F_15_) and three resistant families (F_21_, F_23_, and F_48_) were used as recipients, and 15 families were used as donors. All families were maintained under controlled bio-secured conditions to ensure their specific pathogen-free status. Status was verified by i) the absence of OsHV-1 DNA using qPCR, (see the protocol below) and ii) a low *Vibrio* load (~10 cfu/g tissue) on selective culture medium (thiosulfate-citrate-bile salts-sucrose agar) [24]. Oysters remained free of any abnormal mortality throughout larval development, at which time experimental infections were started.

### Mesocosm experimental infections

The experimental infection protocol consisted of a cohabitation assay between donors (which had been exposed to pathogens naturally present in the environment) and recipient specific pathogen-free oysters [18,19]. Details of the experimental infection protocol (*e.g*., biomass, oyster weight, experimental duration, and tank volume) were as described in [21,23]. Briefly, donor oysters were deployed at Logonna Daoulas (lat. 48.335263, long. −4.317922) in French Brittany (Atlantic environment) and at Thau Lagoon (lat. 43.418736, long. 3.622620) (Mediterranean environment). Both sites differ by a series of ecological and environmental factors but POMS mortalities occur in both sites when temperature reaches ~16°C [25]. Oysters were deployed in farming areas during the infectious period (in July for Atlantic environment and in September for Mediterranean environment, temperature around 21°C for both sites), and remained in place until the onset of donor mortality (< 1%). Donors were then brought back to the laboratory (in Argenton, French Brittany) and placed in tanks, each containing recipient oysters from the three resistant and the three susceptible families. Experimental infections took place in July 2015 and September 2015 for the Atlantic and Mediterranean exposures, respectively. For each experimental infection, mortality rate was monitored, and 10 oysters were sampled in triplicate from each oyster family shucking at 7 time points (0, 6, 12, 24, 48, 60, and 72 hours post-infection). The shell was removed and the whole oyster was flash frozen in liquid nitrogen. Oyster pools (10 oysters per pool) were ground in liquid nitrogen in 50 ml stainless steel bowls with 20mm diameter grinding balls (Retsch MM400 mill). The powders obtained were stored at −80°C prior to RNA and DNA extraction.

### DNA extraction and quantification of OsHV-1 and total bacteria

DNA extraction was performed as described in [21] using the Nucleospin tissue kit (Macherey-Nagel). DNA concentration and purity were checked with a NanoDrop One (Thermo Scientific). Quantification of OsHV-1 and total bacteria were performed using quantitative PCR (qPCR, Roche LightCycler 480 Real-Time thermocycler) with the following program: enzyme activation at 95□ °C for 10□min, followed by 40 cycles of denaturation (95□ °C, 10□s), hybridization (60□°C, 20□Ms) and elongation (72□°C, 25□s). The total qPCR reaction volume was 1.5 □μL with 0.5□ μl of DNA (40□ng□μl-1) and 1□μL of LightCycler 480 SYBR Green I Master mix (Roche) containing 0.5□μM of PCR primers. Absolute quantity of OsHV-1 was determined using virus-specific primer pair targeted the OsHV-1 DNA polymerase catalytic subunit (AY509253, Fw: 5’-ATTGATGATGTGGATAATCTGTG-3’ and Rev: 5’-GGTAAATACCATTGGTCTTGTTCC-3’) and was calculated by comparing the observed Cq values to a standard curve of the DNA polymerase catalytic subunit amplification product cloned into the pCR4-TOPO vector (Invitrogen). Relative quantification of total bacteria 16S rDNA gene was determined using primer pair targeting the variable V3V4 loops (341F: 5’-CCTACGGGNGGCWGCAG-3’ and 805R: 5’-GACTACHVGGGTATCTAATCC-3’) [26] and was calculated by the 2 ^ΛΛCq^ method [27] with the mean of the measured threshold cycle values of two reference genes (*Cg-BPI*, GenBank: AY165040, *Cg-BPI* F: 5’-ACGGTACAGAACGGATCTACG-3’ and *Cg-BPI* R: 5’-AATCGTGGCTGACATCGTAGC-3’ and *Cg-actin*, GenBank: AF026063, *Cg-actin* F: 5’-TCATTGCTCCACCTGAGAGG-3’ and *Cg-actin* R: 5’AGCATTTCCTGTGGACAATGG-3’) [21].

### Analyses of bacterial microbiota

Bacterial metabarcoding was performed using 16S rRNA gene amplicon sequencing. Libraries were generated using the Illumina two-step PCR protocol targeting the V3-V4 region [26]. A total of 252 libraries (six families × seven sampling time points × three replicates × two infectious environments) were paired-end sequenced with a 2 × 250 bp read length at the Genome Quebec platform on a MiSeq system (Illumina) according to the manufacturer’s protocol. A total of 41,012,155 pairs of sequences were obtained. Metabarcoding data was processed using the FROGS pipeline [28]. Briefly, paired reads were merged using FLASH [29]. After cleaning steps and singleton filtering, 26,442,455 sequences were retained for further analyses. After denoising and primer/adapter removal with CUTADAPT, clustering was performed using SWARM, which uses a two-step clustering algorithm with a threshold corresponding to the maximum number of differences between two Operational Taxonomic Units (OTU) (denoising step d = 1; aggregation distance = 3) [30]. Chimeras were removed using VSEARCH [31]. Resulting OTUs were affiliated using Blast+ against the Silva database (release 128).

### Bacterial metatranscriptomic data

Powder obtained from the frozen oysters was resuspended in Trizol, and total RNA was extracted using a Direct-zol^TM^ RNA Miniprep kit. Polyadenylated mRNAs (*i.e*., oyster mRNAs) were removed using a MICROB*Enrich*™ Kit (Ambion). cDNA oriented sequencing libraries were prepared as described in [22] using the Ovation Universal RNA-Seq system (Nugen). Library preparation included steps to remove oyster nuclear, mitochondrial, and ribosomal RNAs, as well as bacterial rRNAs [22]. A total of 36 libraries (three families × two sampling timepoints × three replicates × two infectious environments) were sequenced by the Fasteris company (Switzerland, https://www.fasteris.com) in paired-end mode (2 × 150 bp) on an Illumina HiSeq 3000/4000 to obtain 200-300 million clusters per sample (**Supplementary Table 1**).

Raw Illumina sequencing reads from the resulting 72 fastq files (R1 and R2) were trimmed using Trimmomatic v0.38 (in paired-end mode with no minimum length reads removal). rRNA reads (both eukaryotic and bacterial) were removed using SortmeRNA v2.1b with the rRNA Silva database (release 128) and unmerged, using SortmeRNA ‘unmerge-paired-reads.sh’. At this stage, about 9% of reads were removed, underscoring the efficiency of experimental rRNA removal during library preparation (**Supplementary Figure 1**).

To further enrich for bacterial sequences, unpaired reads were successively mapped by Bowtie2 [32] (very-sensitive-local mode) on a multifasta file containing *Crassostrea gigas* genome sequence v9, complemented by *C. gigas* EST (available from NCBI), and a multifasta file containing the sequences of OsHV-1 (present in diseased oysters) and other viral sequences previously associated with bivalves [33]. Unmapped reads, which represented 4-10% of the starting reads (depending on conditions) were retained for further analysis (**Supplementary Table 1**). Trimmomatic was used again to retrieve paired-reads and remove reads less than 36 nt long. All remaining reads corresponding to the 36 samples were pooled (516,786,580 reads, 36-150 nt) and assembled using Trinity v2.3.2 in paired-end, default mode to build a reference metatranscriptome (1,091,409 contigs, 201-15,917 nt). The resulting metatranscriptome was annotated using Diamond BlastX against the NCBI nr protein database [34]. 48.4 % of the contig encoded proteins aligned with a protein in the database, and were further assigned to a taxa using Megan 6-LR Community Edition [35]. Sequences were annotated at different taxonomic levels from species to phylum. Out of the 1,091,409 contigs, 352,473 contig-encoded ORFs aligned with bacterial proteins by BlastX with an E-value ≤ 01^e^-06 and constituted the bacterial metatranscriptome. For each contig, the best hit was kept. In this metatranscriptome, 54,359 annotated proteins were encoded by the seven genera which were retained for further analysis.

In addition of the gene level, genes were expertly annotated at three functional levels: functions, subcategories and functional categories. First, we defined 31 functional categories (**Supplementary Table 2**). Out of the 54,359 proteins, 9,649 were annotated as “hypothetical”, “unknown”, or “unnamed”, and were assigned to the category “Unknown function”. Using information present in protein databases (NCBI protein, Uniprot, etc.), such as PFAM domains, KEGG number, GO annotation, each unique protein was manually assigned to one of the 30 remaining functional categories. Secondly, each protein was assigned to subcategories, and genes coding for a same function in the same genus (subunits of the same protein complex or enzymatic activity; orthologues with the same function) were manually grouped to produce a reduced table of 9,975 functions.

### Quantification of gene expression and data normalization

For each of the 36 samples used for the assembly of the metatranscriptome, reads were mapped back onto the bacterial metatranscriptome by Bowtie2 in paired-end mode. Raw counts per features (*i.e*., per contig) were computed using HTseq-Count [36]. For each contig, and for each sample, raw counts were normalized to TPM (Transcripts per Kilobase / Million = Mapped reads / Length of contig (kb) / Scaling factor with Scaling factor = total reads in a sample / 1,000,000), which corrects for contig length and differences in read number in the different samples. In many cases, the same protein (having the same unique ID) could be encoded by several contigs, either because gene assembly into a contig was non-contiguous, or because of the existence of contig isoforms.

When analyzing the functions in the seven genera that predominated in the microbiota of diseased oysters, such contigs that encoded the same protein were merged into a single annotation, and their expression levels were summed prior to further normalizaion. First, a pseudo count of one read was added to each gene in each condition/replicate to avoid dividing by zero, then the resulting counts were normalized by dividing by the total number of counts of the genus in a given condition/replicate, and further multiplying by 10,000. For a given gene or function, differential expression was defined as the ratio of the average normalized expression level of the replicates at T60 or 72 over the average normalized expression level at T0, called the expression ratio (ER).

### Statistical analyses

Survival curves were used to determine differential mortality kinetics between oyster families with the non-parametric Kaplan-Meier test (Mantel–Cox log-rank test, p < 0.05, GraphPad_Prism 6.01). For OsHV-1 and total bacteria quantifications, significant differences between resistant and susceptible oyster families were determined using the non-parametric Mann Whitney test (p < 0.05, GraphPad_Prism 6.01). For bacterial metabarcoding, statistical analyses were performed using R v3.3.1 (http://www.R-project.org, [37]). Principal coordinate analysis (PCoA, “phyloseq”) on a Bray-Curtis distance matrix (ordinate, “phyloseq”) was performed to determine dissimilarities between samples. Multivariate homogeneity of group dispersions was tested between bacterial assemblages of the six oyster families using 999 permutations (permutest, betadisper, “vegan”). DESeq2 (“DESeq”, [38]) from OTUs to the higher taxonomic ranks was used to identify candidate taxa whose abundance changed between the initial and final time points of the experiment. Heatmaps of significant genera were computed using relative abundances and the heatmap2 function “ggplots” [39]. For bacterial metatranscriptomics, significance of differential expression between two conditions (*i.e*., T60 or T72 vs T0) was assessed at the level of genes and functions using Student’s t-test (“t.test” function) after controlling for the presence of at least three values (reads in three replicates) in one condition and for variance homogeneity (“var.test” function). Functional enrichment analyses were computed using genes that were significantly differentially expressed to identify over- and underrepresented functional categories or subcategories. These analyses were done for each genus using the list of significant genes (up or down) and the Fisher’s exact test (R package {stats}, fisher.test). P-values of metatranscriptomics were corrected for multiple comparisons using Benjamini and Hochberg’s method (“p.adjust” function) (false discovery rate (FDR) < 0.05).

Lastly, a permutational approach was used to test if the number of specific overexpressed metabolic functions was higher than expected randomly in each environment. This analysis was done on the whole and the core functions (*i.e*., functions shared by the seven genera) in order to test specificity on a similar bacterial genetic background. The significance was assessed by resampling without replacement (“sample” function, MASS package) the metabolic dataset to draw out the expected null distribution. More precisely, we made 999 random matrices of the number of overexpressed functions identified in the seven genera using the reference dataset of each environment. We then compared the observed value to the expected distribution to compute a p-value based on the number of random samples that showed higher number of specific functions.

### Data availability

Metabarcoding and RNAseq sequence data are available through the SRA database (BioProject accession number PRJNA423079). For bacterial metatranscriptomic, SRA accessions of BioSamples were SAMN15461557 to SAMN15461592. For bacterial metabarcoding SRA accessions of BioSamples were SAMN15462520 to SAMN15462771. Large supplementary files are available in the OSF database (https://osf.io/kybva/). Other data generated from this study are included in the published version of this article and its supplementary files.

## RESULTS

### Primary OsHV-1 infection and secondary bacteremia are conserved in POMS, independently of the infectious environment

Six *C. gigas* families were subjected to two experimental infections mimicking disease transmission in the wild. We previously reported high variability in the dynamics of mortality and final percentage survival of oyster families confronted with an Atlantic infectious environment. Specifically, the F11, F14, and F15 families were highly susceptible (survival rate < 4% after 330h) to POMS, whereas the F21, F23, and F48 families were highly resistant (survival rate > 82% after 330h) [21]. Similar results were obtained in the present study when the same oyster families were confronted with a Mediterranean infectious environment: families F11, F14, and F15 were susceptible (survival rates < 9%), whereas families F21, F23, and F48 were resistant (survival rates > 88%) (**Figure 1**). Thus, these oyster families displayed similar phenotypes when confronted with two different infectious environments (Mantel-Cox log-rank test, p < 0.0001 for each comparison of resistant *vs*. susceptible oyster families). Susceptible and resistant oyster families are hereafter referred to as S (S_F11_, S_F14_, and S_F15_) and R (R_F21_, R_F23_, and R_F48_), respectively.

**Figure 1.**
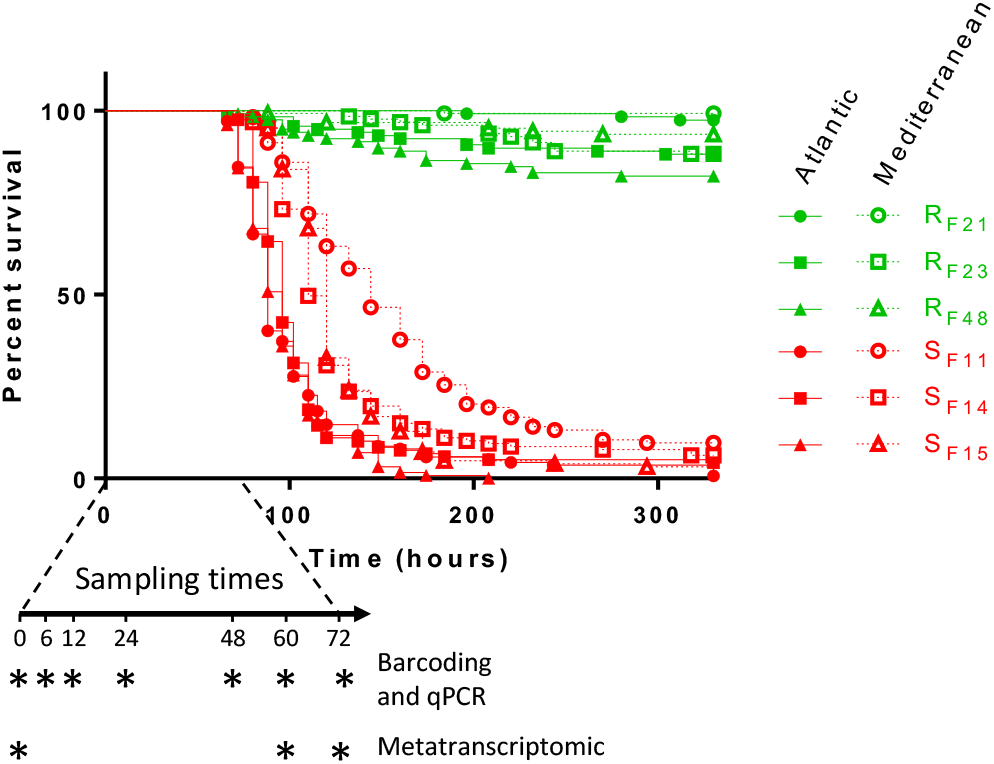
Kaplan-Meier survival curves of oyster biparental families confronted with two different infectious environments. Resistant oyster families (R_F21_, R_F23_, and R_F48_) are presented in green, and susceptible oyster families (S_F11_, S_F14_, and S_F15_) are presented in red. At each time point (indicated by asterisks on the arrow), 10 oysters were sampled 3 triplicates from each family in each tank for barcoding, qPCR, and metatranscriptomic analysis. For metatranscriptomic analysis oysters were sampled at the onset of mortalities (60h and 72h post-exposure for Atlantic and Mediterranean infectious environments, respectively). Data for the Atlantic infectious environment was extracted from [21] and shown for comparison.

We then compared pathogenesis between the two infectious environments by monitoring OsHV-1 load, microbiota dynamics, and bacterial abundance in the three resistant and three susceptible oyster families (**Figure 2**). OsHV-1 DNA was detected in all families, regardless of whether they were confronted with the Atlantic or Mediterranean infectious environment (**Figure 2a)**. However, very intense viral replication occurred only in the susceptible oyster families: viral DNA loads were 2 to 3 logs higher than in resistant oysters at 24 h (**Figure 2a**).

**Figure 2.**
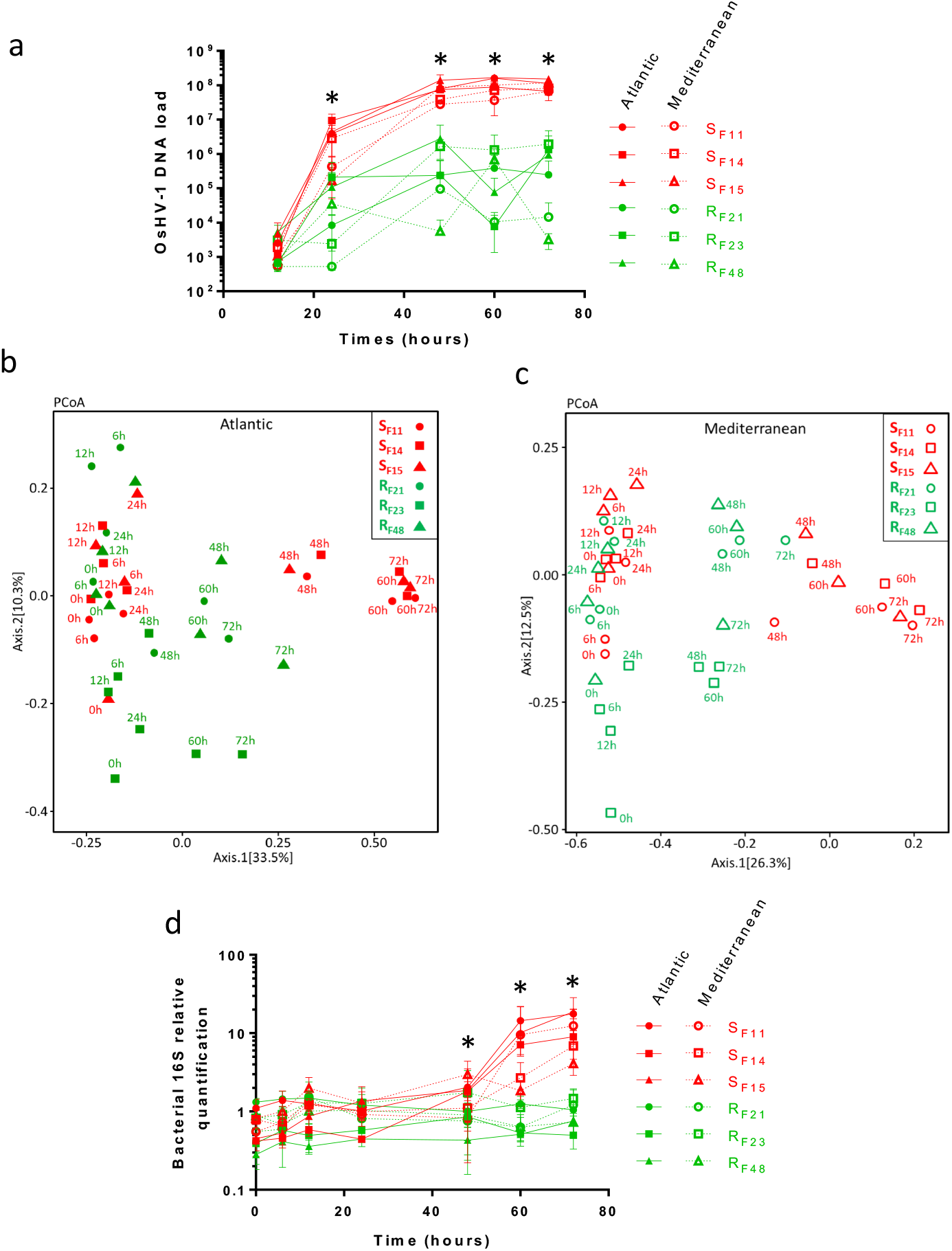
Primary OsHV-1 infection, bacterial dysbiosis, and secondary bacteremia are conserved in different infectious environments. **(a)** Early and intense replication of OsHV-1 μVar occurs in susceptible oysters (red), but not resistant oysters (green), confronted with either the Atlantic or the Mediterranean infectious environment. OsHV-1 load was quantified by qPCR and expressed as Viral Genomic Units per ng of oyster DNA (log scale) during experimental infections. Asterisks indicate significant differences between susceptible and resistant oyster families (Mann Whitney test, p < 0.05). **(b-c)** Principal coordinate analysis (PCoA) plot of the microbiota for susceptible (red) and resistant (green) oyster families confronted with each infectious environment. Dispersion of oyster families according to the Bray-Curtis dissimilarity matrix (beta diversity) in **(b)** Atlantic and **(c)** Mediterranean infectious environments. **(d)** Temporal dynamics of total bacteria in susceptible (red) and resistant (green) oyster families confronted with two different infectious environments. Total bacterial quantification based on qPCR amplification of the V3-V4 region of the 16S rRNA gene during experimental infections. Asterisks indicate significant differences between susceptible and resistant oyster families (Mann Whitney test, p < 0.05). Data from the Atlantic infectious environment in panels (a) and (d) are extracted from [21] for comparison.

The dynamics of the oyster microbiota was studied in the six oyster families by monitoring bacterial community composition using 16S rRNA gene metabarcoding over the first 3 days of both experimental infections. A total of 45,686 bacterial OTUs were obtained from the 252 samples and affiliated at different taxonomic ranks (**Supplementary Table 3**). Changes in microbiota composition were greater in susceptible oysters than in resistant oysters at all taxonomic ranks (**Supplementary Figure 2**). Indeed, for the Atlantic infectious environment, 52, 43, and 54 OTUs significantly differed (in terms of relative abundance between the start and end of the experiment) in susceptible oysters S_F11_, S_F14_ and S_F15_, respectively; only 1, 11, and 9 OTUs significantly differed in resistant oysters R_F21_, R_F23_ and R_F48_, respectively (**Supplementary Table 4**). The same trend was observed in the Mediterranean infectious environment. 11, 47, and 43 OTUs significantly differed in S_F11_, S_F14_ and S_F15_, respectively, as opposed to 2, 8, and 6 OTUs in R_F21_, R_F23_ and R_F48_, respectively.

PCoA on a Bray-Curtis dissimilarity matrix (beta diversity) revealed higher microbiota dispersion in susceptible oyster families than in resistant families in both infectious environments (multivariate homogeneity of groups dispersion, d.f. = 1; *p* = 0.016 and *p* = 0.020 for Atlantic and Mediterranean environments, respectively) (**Figure 2b and 2c**). This disruption of the bacterial community structure occurred in susceptible oysters between 24 h and 48 h, concomitantly with the active replication of OsHV-1. In addition, susceptible oyster families displayed a significantly greater bacterial load than resistant oysters when confronted with either the Atlantic or the Mediterranean infectious environment (Mann Whitney test, p < 0.05; **Figure 2d**). This increase started at 60 h and continued until the end of the experiment (72 h). Total bacterial abundance in susceptible oysters was more than 5-fold higher at 72 h than at T0, which indicated bacterial proliferation. In contrast, total bacterial load remained stable in resistant oysters.

### A core pathobiota infects oysters during secondary bacterial infection in POMS

All bacterial genera that changed significantly in abundance during the two experimental infections (Atlantic and Mediterranean) in susceptible oyster families are reported in **Supplementary Table 4.** We focused on well-represented genera representing > 2% of the bacteria in at least one sample for each susceptible oyster family confronted with each infectious environment (**Figure 3)**. In the Atlantic infectious environment and for susceptible families, the corresponding OTUs represented a total of 4%, 0.8%, and 46% of total bacteria at the beginning of the experiment (T0), as opposed to 73%, 75%, and 72% at 72 h for S_F11_, S_F14_, and S_F15_, respectively (**Supplementary Table 4**). In the Mediterranean infectious environment and for susceptible families, these OTUs increased from 2%, 6%, and 7% at T0 to 47%, 56%, and 56% at 72 h for S_F11_, S_F14_, and S_F15_, respectively. From nine to twenty genera increased significantly in relative abundance between T0 and 72 h. Ten genera (*Arcobacter, Cryomorphaceae, Marinobacterium, Marinomonas, Proxilibacter, Pseudoalteromonas, Psychrilyobacter, Psychrobium, Psychromonas*, and *Vibrio*) were common to almost all (5 of 6) susceptible oyster families and both infectious environments (**Figure 3**). Most of the remaining genera (*Aquibacter, Aureivirga, Fusibacter, Neptunibacter, Peredibacter, Pseudofulvibacter*) were shared by at least two families in one infectious environment. One genus (*Salinirepens*) increased significantly in all susceptible oysters in the Atlantic infectious environment only. These results show that a core of bacterial genera infects oysters during the POMS secondary bacterial infection, independently of the infectious environment. In resistant oyster families, several taxa also varied significantly in abundance over time. Most of these taxa were also present in susceptible oyster families (**Supplementary Figure 3)**, but at low abundances. These taxa represent between 4% to 23% of the reads sequenced at 72h in resistant oysters, whereas they represent 47% to 75% of the reads sequenced in susceptible oysters (**Supplementary Table 4**).

**Figure 3.**
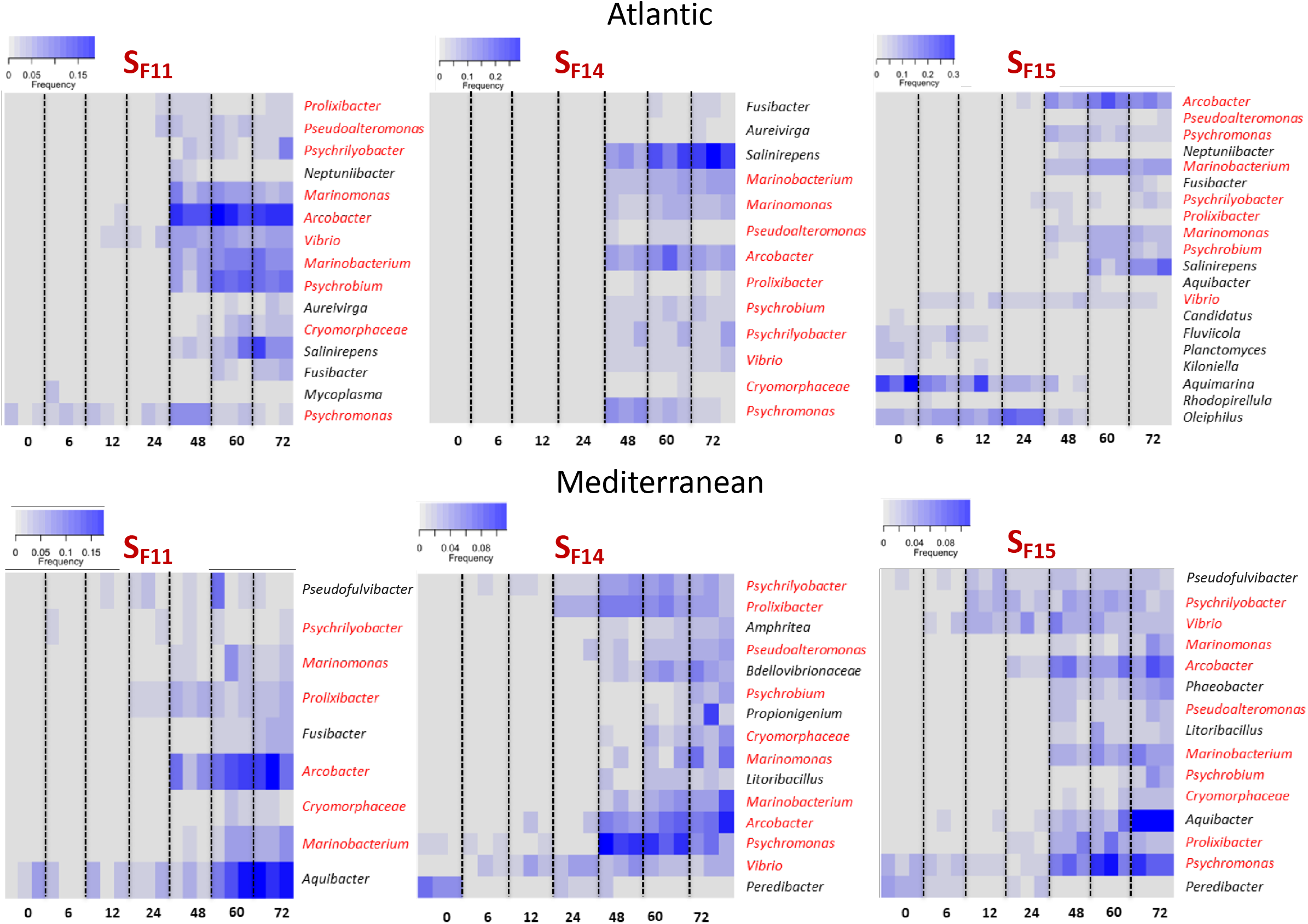
Heatmaps of bacterial genera that changed significantly in abundance over the course of infection in susceptible oysters (S_F11_, S_F14,_ and S_F15_) in the Atlantic and Mediterranean infectious environments. Analyses were performed at the genus level. Only genera that changed significantly in abundance and had a relative proportion greater than 2% in at least one sample are shown. Increased intensity of color (blue) represents increased relative abundance. Genera that are consistently modified in 5 out of the 6 conditions (3 families and 2 infectious environments) are in red.

### Seven genera are responsible for most bacterial gene expression in diseased oysters

To understand the infection success of certain genera, we analyzed the gene expression of the pathobiota using metatranscriptomics. As the secondary bacterial infection did not occur in resistant oysters, it seemed difficult to obtain from these oysters a sequencing depth for bacteria sufficient for subsequent analysis, and we chose to restrict the metatranscriptomic analysis to the three different susceptible families S_F11_, S_F14_, and S_F15_, from both Atlantic and Mediterranean infectious environments, at T0 and just before oyster mortality occurred (*i.e*., at 60 h and 72 h for the Atlantic and the Mediterranean infectious environments, respectively). Three biological replicates were analyzed for each condition, corresponding to a total of 36 biological samples, 8,4 billion reads assembled into 352,473 contigs, and 225,965 unique proteins.

Seven genera were consistently found to contribute to most of transcriptomic activity in diseased oysters, displaying a strong relative increase of the number of transcripts compared to healthy oysters (**Figure 4**). *Amphritea, Arcobacter, Marinobacterium, Marinomonas, Oceanospirillum, Pseudoalteromonas*, and *Vibrio* were together responsible for up to 40% of the total bacterial transcriptomic activity detected just before the onset of oyster mortality. Among them, only *Amphritea* and *Oceanospirillum* were not part of the core pathobiota identified using metabarcoding, even though *Amphritea* was significantly increased in S_F14_. Six out of seven genera are Gammaproteobacteria: while *Amphritea, Marinobacterium, Marinomonas* and *Oceanospirillum* belong to the same family (Oceanospirillaceae) and order (Oceanospirillales), *Arcobacter* belongs to the class Epsilonproteobacteria.

**Figure 4.**
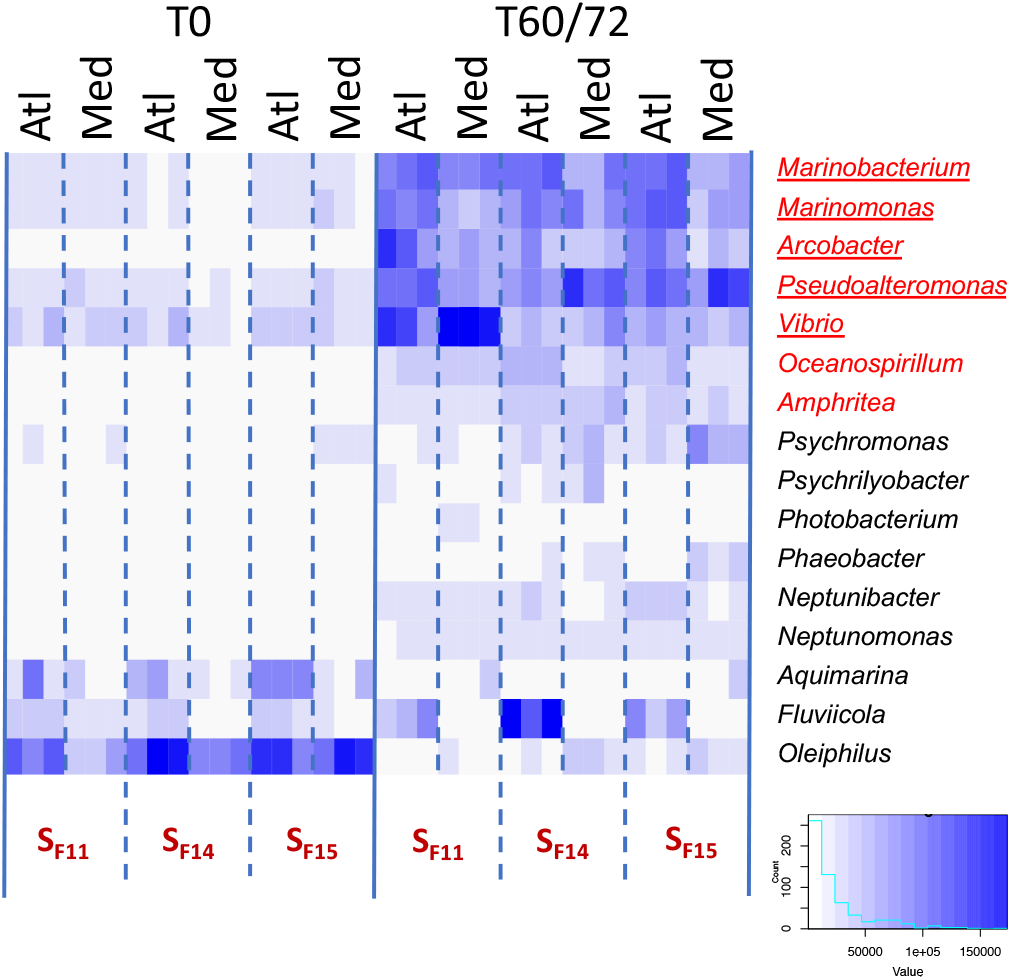
Heatmap of transcriptional activity of bacterial genera in susceptible oyster families (S_F11_, S_F14_, and S_F15_) in the two infectious environments at the time of exposure to the infectious environment and in diseased oysters. For each condition, results of the three replicates are shown. Increased color intensity (blue) indicates increased relative activity of the genus. Genera shown contributed at least 2% of the total transcriptional activity in at least one sample of diseased oysters. Bacterial genera that were overrepresented according to metatranscriptomics alone for all conditions in the diseased oysters are in red, while genera that were overrepresented according to both metabarcoding and metatranscriptomics are underscored (Atl: Atlantic, Med: Mediterranean, T0: time of exposure to the infectious environment, T60/72: diseased).

These results indicate that a limited number of genera participate in the secondary bacteremia that occurs in POMS. These genera are remarkably conserved between the different susceptible oyster families. Therefore, we focused our analyses on these seven genera, considering samples from all three susceptible families as replicates and comparing two time points (T0 *vs*. 60 h or 72 h for the Atlantic or the Mediterranean infectious environments, respectively) and the two different environments. The pathobiota constituted by these seven genera corresponded to 106,312 contigs and 54,359 unique proteins (query and subject in **Supplementary Table 5**).

### The seven bacterial genera showed reproducible differential expression patterns in both environments

For each genus, and each infectious environment, normalized expression levels were estimated at the gene level (**Supplementary Table 6**). To each gene was attributed a function, a functional category and a subcategory.

We first compared the variation of the expression pattern at the level of functional categories between the time of exposure to the infectious environment (T0) and the onset of mortality (*i.e*., at 60 h and 72 h for the Atlantic and the Mediterranean infectious environments, respectively), for the seven bacterial genera in both environments (**Figure 5**). With the exception of the category “Translation/ ribosomal structure and biogenesis” (translation for short), which was globally overexpressed, a striking fact was the specific differential expression pattern of each genus. A clustering analysis of the differential expression of the functional categories showed that, for a given genus, the two environments generated a very similar pattern of differential gene expression.

**Figure 5.**
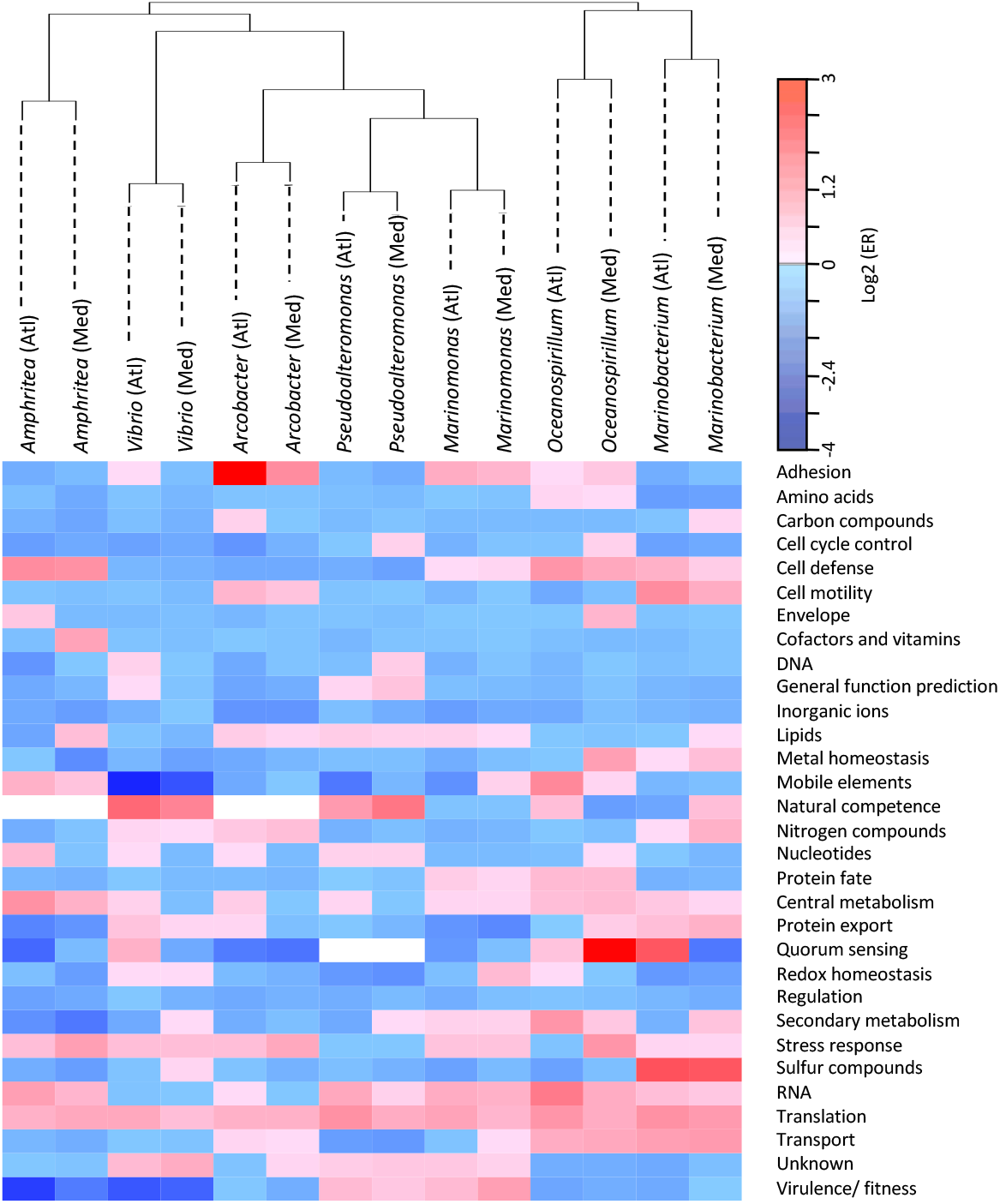
Heatmap of bacterial gene expression variation for each 31 functional categories between T0 and the onset of oyster mortality. Graded colors (blue to red from decreases to increases) are used to represent the extent of the global changes of each category, using a log 2 scale. White cells indicate categories with no gene expression in a given genus (gene absent or expression not detected).

### Translation and central metabolism were significantly overexpressed by pathobiota during the infectious process

To identify the genes underpinning successful infection by the seven bacterial genera, we next computed a comparison between T0 and the onset of mortality. For each genus, and each infectious environment, expression levels were compared at the gene level (**Supplementary Table 6**). For each bacterial genus, a majority of genes were overexpressed within each environment (**Supplementary Figure 4**), and overexpression seemed to affect more genes in the Atlantic than in the Mediterranean environment.

Gene set enrichment analyses were carried-out using significantly over- or underexpressed genes from the **Supplementary Table 6** to identify over- and underrepresented functional categories. While no category was significantly enriched for underexpressed genes, the translation category was found to be overrepresented in the overexpressed genes, for all genera and in both environments (except *Vibrio* in the Mediterranean environment) (**Supplementary Figure 5**), consistent with a general increase of expression of genes from this category in diseased oysters (**Figure 5**). In addition, the category “Precursor metabolite and energy production” (central metabolism for short) was significantly enriched in the overexpressed genes of four out of seven genera, *Amphritea, Marinobacterium, Marinomonas* and *Oceanospirillum*.

We then computed enrichment analyses within both categories of translation and central metabolism in order to identify overrepresented subcategories. The two subcategories of 30S and 50S ribosomal proteins were found to be overrepresented for most genera amongst the overexpressed genes (**Supplementary Figure 6**) whereas oxidative phosphorylation was overrepresented for two genera in the case of the central metabolism category (**Supplementary Figure 7**).

### Metabolic complementarity might explain reproducible composition of bacterial assemblages

The enrichment during the infection process in the overexpressed gene set of the central metabolism category (**Supplementary Figure 5**), and, especially, of the oxidative phosphorylation subcategory, (**Supplementary Figure 7**) suggest that changes in metabolic activity are important for the successful establishment of the pathobiota.

Accordingly, we focused the next analyses on genes involved in metabolic categories. In particular, we compared the different genera for the differential expression of transcripts from the categories of amino acids, carbon compounds, lipids, nitrogen compounds, central metabolism, and sulfur compounds (see **Supplementary Table 2**). To perform the analyses at the scale of metabolic functions, the whole dataset was reduced by grouping genes having the same function (subunits of the same protein or enzymatic complex; see Methods). Then, for each genus and each infectious environment, expression levels were compared between T0 and the onset of mortality (*i.e*., at 60 h and 72 h for the Atlantic and the Mediterranean infectious environments, respectively) for all functions (**Supplementary Table 7**).

The seven genera showed very few similarities in terms of significantly overexpressed metabolic functions. Indeed, only 5 out of 222 metabolic functions were increased in at least four genera in the Atlantic or the Mediterranean environment, all of them belonging to the category of central metabolism: ATP synthase (oxidative phosphorylation), dihydrolipoyl dehydrogenase (pyruvate metabolism), cytochrome-c oxidase, cbb3-type (respiratory electron transfer), glyceraldehyde 3-phosphate dehydrogenase and triose-phosphate isomerase (glycolysis/gluconeogenesis) (**Supplementary Table 8**).

Most overexpressed metabolic functions were specific of a single genus in the Atlantic (68.39%; 119/174 functions) and in the Mediterranean environment (77.08%; 37/48). In order to estimate the significance of these specificities, permutational analyses were computed and revealed that these high ratios of specific functions were higher than expected randomly (p=0.001 in both environments). These analyses were also done on the core metabolic functions (*i.e*., functions shared by the seven genera) in order to avoid bias due to different genetic backgrounds. For the overexpressed core functions, 54.55% (36/66) and 72% (18/25) were specific of a single genus in the Atlantic and the Mediterranean environment, respectively. Permutational analyses also highlighted that these high ratios were significant (p=0.001 in both environments).

### Oysters provide a diverse set of nutritive sources to the pathobiota

In addition to **Supplementary Table 8**, **Figure 6** and **Supplementary Figure 8** present a schematic view of the main metabolic changes in the pathobiota and the respective contribution of each genus. All of them contributed to some extent to an increase of the main pathways of central metabolism, glycolysis/neoglucogenesis, ß-oxidation, TCA cycle, respiratory electron transfer and oxidative phosphorylation, and pentose phosphate and PRPP biosynthesis but each of them in a specific way. Overall, the pathobiota metabolic network reflected the diversity of the nutrients available in the diseased oysters. Thus, amino acids, in degraded tissues, can be a major source of carbon, neoglucogenesis being favored to glycolysis in central metabolism, as seen in *Marinobacterium, Oceanospirillum, Pseudoalteromonas*, and *Vibrio*. Consistent with this was the increase of several amino acid degradation pathways in these genera. In this rich environment, other carbon sources are available: aromatic compounds and xenobiotic compounds such as atrazine (*Marinomonas*) or isoprene (*Amphritea, Pseudoalteromonas*). Glycogen, which is especially abundant in oysters, and host glycans, are potential sources of glucose. *Pseudoalteromonas* and *Vibrio* showed an increase in the use of several sugars and sugar derivatives (**Supplementary Table 8**).

**Figure 6.**
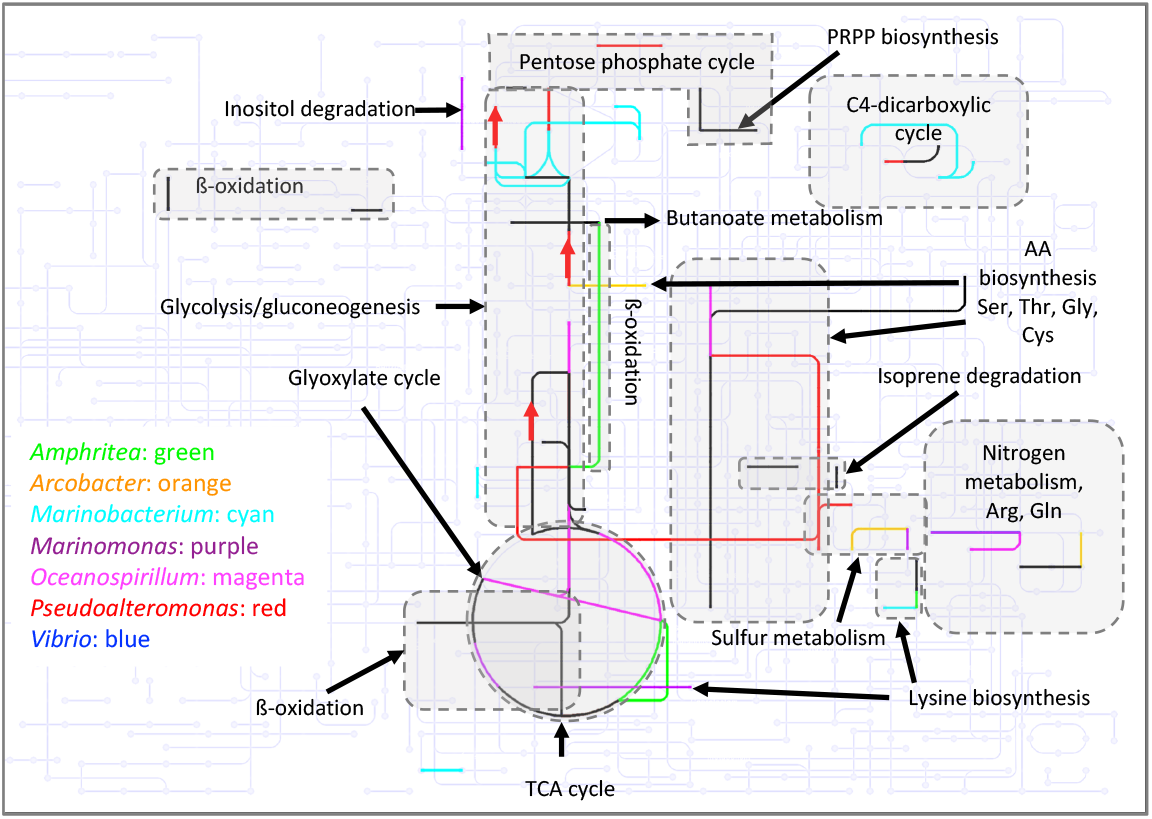
Increased expression pathways of the pathobiota metabolism between T0 and the onset of oyster mortality and the contribution of each bacterial genus. EC numbers of differentially overexpressed genes involved in bacterial metabolism **(Supplementary Table 8**) were mapped on KEGG metabolic map 01120 (Microbial metabolism in diverse environments, https://www.kegg.jp/kegg-bin/show_pathway?ec01120) using a color code for each genus. The highlighted pathways are labelled. Pathways common to two genera or more are in black. Red arrows indicate the pathway corresponding to neoglucogenesis. Note that not all relevant pathways are represented on this map (such as oxidative phosphorylation) which was chosen for the sake of clarity.

In dying oysters, the environment could evolve towards microaerobic to anaerobic conditions, favoring in certain genera the activation of nitrate respiration (*Amphritea, Arcobacter, Marinobacterium* and *Marinomonas*) or L-carnitine respiration (*Marinomonas*, Atlantic environment). In addition, the formate dehydrogenase (increasing in *Arcobacter* and *Oceanospirillum*) could also play a role in nitrate or other final electron acceptor respiration. Oysters are also rich in taurine [40], whose catabolism is highly induced in *Marinomonas*. In addition to carbon, and nitrogen, Taurine can also be a source of sulfur.

Changes of expression of several sulfur metabolism pathways were also observed in all genera, except in *Oceanospirillum* and *Vibrio*. Finally, activation of ß-oxidation (*Amphritea, Marinobacterium* and *Pseudoalteromonas*) indicates that fatty acids are also used as nutrients.

### Specific patterns of adaptive responses in the different bacterial genera during the infection

Other functions that might be key to successful host infection are functional categories that are important for survival in the host and/or pathogenicity, such as adhesion, cell defense, metal homeostasis, redox homeostasis and oxidative stress, stress response, and virulence factors (**Supplementary Table 9**). Each genus displayed some responses to such stresses, with varying specific strategies. We found a varying repertoire of genes involved in the maintenance of intracellular reducing potential, such as thioredoxin and glutaredoxin, that play an important role in reducing protein disulfide bonds in the cytoplasm [41] as well as genes coding for peroxidase and superoxide dismutases, which are important for the oxidative stress response, whose expression was either decreased or increased. For metal homeostasis, the main response was to maintain iron concentration. More widely shared between genera, was the induction of genes coding for formaldehyde dehydrogenase, an aldehyde-detoxifying enzyme, of cold shock protein genes and genes coding for the ribosome-associated translation inhibitor RaiA. Interestingly, cold shock proteins are often RNA chaperons which are important for ribosome biogenesis [42]. The induction of RaiA and cold shock proteins could reflect the very high translation activity in the pathobiota. Finally, in the fitness/virulence gene category, genes whose expression was significantly affected belonged to the fitness (competition between bacteria) rather than host interactions category.

## DISCUSSION

### POMS pathobiota is composed of a few and reproducible number of active bacterial genera

Until recently, only members of the *Vibrio* genus had been repeatedly associated with POMS. These studies used culture-based approaches to investigate oyster-associated bacterial communities [19,20]. *Vibrio* species associated with POMS were characterized by key virulence factors that are required to weaken oyster cellular defenses [22,43]. Members of the *Arcobacter* genus had also been associated with POMS-diseased oysters [44,45], but the role of this genus in pathogenesis was not investigated more deeply due to limitations of culture-based techniques [46]. In the present study, we showed that the dysbiosis associated with POMS was conserved across infectious environments. Using metabarcoding, we demonstrated that diseased oysters affected by POMS are colonized by a common consortium of bacteria composed by ten major genera (*Arcobacter, Cryomorphaceae, Marinobacterium, Marinomonas, Proxilibacter, Pseudoalteromonas, Psychrilyobacter, Psychrobium, Psychromonas*, and *Vibrio*), whereas metatranscriptomic data showed that five of these genera (*Arcobacter, Marinobacterium, Marinomonas, Pseudoalteromonas* and *Vibrio*) displayed a high transcriptomic activity, and identified two additional active genera (*Amphritea* and *Oceanospirillum*), thus extending the core bacterial consortium to five additional bacterial genera. The discovery of the contributions of these genera, which are responsible for up to 40% of the bacterial transcriptional activity observed in POMS, provides new insights into the pathogenesis. Altogether, our results strongly suggest that a core microbiota, rather than specific bacterial pathogens, operates as a functional unit of pathogenesis. Together with OsHV-1, these bacteria form the POMS pathobiota. POMS secondary bacteremia may resemble periodontitis in humans, in which the evolution of the disease is characterized by the development of a pathogenic consortium comprising a limited number of species [47,48].

We used metatranscriptomics to unveil the functions of the microbiota in relation to POMS. Bacterial metatranscriptomics from host tissues is technically challenging (due to the low proportion of bacterial transcripts in the host samples), but it provides functional information that is thought to more accurately portray the role of the microbiota in health and disease states [49]. Accordingly, gene expression profiling has proven highly successful in advancing the understanding of the dynamics of disease-associated microbial populations [50]. In the case of POMS, by linking functional genes to the bacterial genera which encode them, we found a remarkably consistent relationship between the structure of bacterial communities (using 16S metabarcoding) and the functions expressed by bacterial genera in the communities (using metatranscriptomics), with the exception of *Amphritea* and *Oceanospirillum*, which were not detected as significantly more abundant at the onset of mortality by metabarcoding (except *Amphritea* in one condition) despite their significant contribution to the pathobiota transcriptional activity. This might be explained by the fact that detection of transcriptional activity might be more sensitive than metabarcoding, allowing detection at an earlier time [51], suggesting that these two genera might become abundant at a later step of oyster infection.

### A pathogenicity independent of bacterial virulence factors?

Surprisingly, only a very few numbers of putative virulence genes have been identified as significantly overexpressed in the seven genera. Indeed, overexpressions were significant for a limited number of genes in *Marinomonas* (1 out of 11 genes in the Mediterranean environment), and *Pseudoalteromonas* (4 and 3 out of 76 genes in Atlantic and the Mediterranean environment, respectively). First, this result might be explained by a lack of knowledge concerning virulence genes in the seven genera. Indeed, while more than 76 genes were listed in this category for *Pseudoalteromonas* or *Vibrio* in this study, less than 13 genes were annotated as virulence/fitness for the other genera. Other candidates may be in the unknown function category. However, virulence genes of *Vibrio* are well described [22,43], and none of these genes were significantly modified. It is also possible that virulence genes were only overexpressed at the onset of the infection process, and not anymore at this late stage. Future studies analyzing the microbiota transcriptomes over time could help resolve this infection-related question. In addition, a transcriptomic analysis of oysters will also be useful to describe host responses to the infection and their immune status [21,22].

### Functional reprogramming centered on bacterial metabolism

Our metatranscriptomic analysis highlighted the specificity of the genera that compose the pathobiota, both in term of metabolism expression (**Supplementary Table 8**), and of adaptation to the host (**Supplementary Table 9**). However, a few core functions, that were overexpressed in at least four genera, was also identified. The most conserved response was a strong induction of genes involved in translation (**Supplementary Figures 4 and 5**), constituting a set of genes enriched in ribosomal proteins (**Supplementary Figure 6**). Interestingly, genes coding for cold shock proteins, which are often RNA chaperones involved in translation and ribosome biogenesis [42] were also part of this functional core genes. Finally, genes for “ATP synthase” and “cytochrome c-oxidase, cbb3 type”, encoding two major components of oxidative phosphorylation and respiration, were induced in five and four genera, respectively.

Beside this limited core response, a high and significant proportion of metabolic functions was overexpressed in only one genus in both environments (**Supplementary Table 8**), suggesting that each genus used different sources and different metabolic pathways. Thus, the pathobiota metabolism reflects on one hand the environment provided by immunocompromised and dying oysters (a rich medium, constituting an abundant source of amino acids and lipids, sustaining a high central metabolism and growth rate) and, on the other hand, the specific metabolic expression of each genus.

This specificity might be the basis of a functional complementation between the bacterial genera. First, this complementarity might be the result of synergy between genera through involvement in different steps of biogeochemical cycles; the growth of one genus favoring the growth of others. For example, metabolic complementarity was proposed between two bacterial symbionts of sharpshooters for histidine and essential amino acids based on genomic analyses [52]. However, we did not identify a complementarity similar to a codependency here. In contrast, this complementarity might be linked to low competition for resources between the different genera, suggesting an optimal use of the diversity of resources in the oyster environment, which can sustain efficient growth of bacteria with very different metabolisms. Such a pattern of coexistence through low nutritive competition (also named resource partitioning) was already observed for several taxa, such as fishes [53], hoverflies [54], and honey bee gut bacteria [55]. In this last study, it was demonstrated thanks to metatranscriptomics and metabolomics that bacterial species used different carbohydrate substrates. This result indicated resource partitioning as the basis of coexistence within honey bee gut, and the longstanding association with their host. For oysters, we also hypothesized that the metabolic complementarity identified here using metatranscriptomics might reflect resource partitioning. This complementarity might explain the reproducible nature of pathobiota assemblages associated with POMS across distinct environments.

## Conclusions

Using metabarcoding and metatranscriptomics, we found that seven bacterial genera were consistently present and active in susceptible oysters affected by POMS in two infectious environments. Moreover, we also found a reproducible nature of the pathobiota composition and transcriptional activity between both environments (Atlantic and Mediterranean). Thanks to metatranscriptomics, we proposed that the conservation of this assemblage might be explained by complementary use of resources with lack of competition between genera. Indeed, oyster tissues might offer conserved ecological niches to the pathobiota during infection process in both environments. Future studies should perform metabolic studies of these genera to validate our observations done at the level of gene expression. Lastly, a temporal analysis of gene expressions of both oysters and microbiota will also help understanding this polymicrobial process at the early steps of infection.

## Acknowledgements

We warmly thank the staff of the Ifremer stations of Argenton (LPI, PFOM) and Sète (LER), and the Comité Régional de Conchyliculture de Méditerranée (CRCM) for technical support in the collection of the oyster genitors and reproduction of the oysters. The authors are grateful to Philippe Clair from the qPHD platform/Montpellier genomix for useful advice and to Ifremer bioinformatics department (SeBIMER) for SRA submission. This work benefited from the support of the National Research Agency under the “Investissements d’avenir” program (reference ANR-10-LABX-04-01) through use of the GENSEQ platform (http://www.labex-cemeb.org/fr/genomique-environnementale-2) from the LabEx CeMEB. The present study was supported by the ANR projects DECIPHER (ANR-14-CE19-0023) and DECICOMP (ANR-19-CE20-0004), and by Ifremer, CNRS, Université de Montpellier and Université de Perpignan *via* Domitia. Xing Luo was a recipient of a fellowship from the China Scholarship Council (CSC). This study is set within the framework of the “Laboratoires d’Excellence (LABEX)” TULIP (ANR□10□LABX□41).

## Author contributions

J.D.L., B.P., A.J. and G.M. designed experiments. B.P., J.D.L, A.L., J.M.E., Y.G., L.D. and G.M. performed oyster experiments. J.D.L., A.L., E.T., C.C. and G.M. performed microbiota analyses. J.D.L. and A.L. performed qPCR analyses. A.J. and X.L performed the metatranscriptomic experiments. A.J. C.C. and S.M. analyzed the metatranscriptomic data. J.D.L., A.L., A.J, C.C., S.M. and G.M. interpreted results. J.D.L., A.L., A.J., C.C., D.D.G. and G.M. wrote the manuscript, which has been reviewed and approved by all authors.

## Conflict of interest statement

There are no conflicts of interest. This manuscript represents original results and has not been submitted elsewhere for publication.

## Supplementary data

**Supplementary Figure 1. Steps of metatranscriptomic analyses.**

**Supplementary Figure 2. Microbiota modification as analysed using 16S rRNA metabarcoding in susceptible and resistant oyster families confronted with two different infectious environments.** Susceptible oyster families (S_F11_, S_F14_ and S_F15_) and resistant oyster families (R_F21_, R_F23_ and R_F48_) confronted with (**a**) Atlantic or (**b**) Mediterranean infectious environments. Significant changes in abundance (up or down; DESeq2, FDR < 0.05) between the initial and the final time point of the infection were much greater for each taxonomic rank (from the phylum to the OTU rank) for susceptible oyster families than for resistant oyster families. Data for AS_F11_ and AR_F21_ were extracted from [21].

**Supplementary Figure 3. Heatmaps of bacterial genera that changed significantly in abundance over the course of infection in resistant oysters (R_F11_, R_F14_, R_F15_) in the Atlantic and Mediterranean infectious environments.** Analyses were performed at the genus level. Only genera with a relative proportion greater than 2% in at least one sample are shown. Increased color intensity (blue) indicates increased relative abundance of the genus.

**Supplementary Figure 4. Number of significant over- and underexpressed genes in each genus and each infectious environment.**

**Supplementary Figure 5. Enrichment analysis of significant overexpressed bacterial genes in the 31 functional categories.** Graded colors (blue to red) are used to represent the observed over expected values (Fisher’s exact tests), and indicate under- to overrepresentation, respectively. Grey cells indicate not significant categories.

**Supplementary Figure 6. Enrichment analysis of significant overexpressed bacterial genes within the functional category of translation.** Graded colors (blue to red) are used to represent the observed over expected values (Fisher’s exact tests), and indicate under- to overrepresentation, respectively. Grey cells indicate not significant subcategories.

**Supplementary Figure 7. Enrichment analysis of significant overexpressed bacterial genes within the functional category of central metabolism.** Graded colors (blue to red) are used to represent the observed over expected values (Fisher’s exact tests), and indicate under- to overrepresentation, respectively. Grey cells indicate not significant subcategories.

**Supplementary Figure 8. Decreased expression pathways of the pathobiota metabolism between T0 and the onset of oyster mortality and the contribution of each bacterial genus.** EC numbers of differentially underexpressed genes involved in bacterial metabolism **(Supplementary Table 8**) were mapped on KEGG metabolic map 01120 (Microbial metabolism in diverse environments, https://www.kegg.jp/kegg-bin/show_pathway?ec01120) using a color code for each genus. The highlighted pathways are labelled. Pathways common to two genera or more are in black. Red arrows indicate the pathway corresponding to neoglucogenesis. Note that not all relevant pathways are represented on this map (such as oxidative phosphorylation) which was chosen for the sake of clarity.

**Supplementary Table 1. Total raw reads (R1+R2) at each stage of biocomputing, after sequencing and removal of rRNA reads (eukaryotic and bacterial), oyster reads, and viral reads.**

**Supplementary Table 2. List of functional categories defined for bacterial metatranscriptomics.**

**Supplementary Table 3. Absolute abundance of bacteria and their corresponding taxonomic affiliations in susceptible and resistant oyster families confronted with two different infectious environments.** Susceptible oyster families are S_F11_, S_F14_, and S_F15_; resistant oyster families are R_F21_, R_F23_, and R_F48_. A indicates the Atlantic infectious environment, M the Mediterranean infectious environment. T0, T6, T12, T24, T48, T60, and T72 indicate sampling times (in hours) over the course of experimental infection. R1, R2, R3 indicate the results of each replicate. This large table is available at https://osf.io/kybva/.

**Supplementary Table 4. Frequencies of bacterial taxa that change significantly in abundance over the course of each experimental infection (Atlantic or Mediterranean) in susceptible and resistant oyster families.** The change in abundance of bacterial taxa between initial and final time points was determined using DEseq2 with the FDR < 0.05. Susceptible oyster families are S_F11_, S_F14_, and S_F15_; resistant oyster families are R_F21_, R_F23_, and R_F48_. A indicates the Atlantic infectious environment, M the Mediterranean infectious environment. T0, T6, T12, T24, T48, T60 and T72 indicate sampling times (in hours) over the course of experimental infection. R1, R2, R3 indicate the results of each replicate.

**Supplementary Table 5. Contigs identified from the seven main bacterial genera.** Annotations and expression values in TPM are indicated. This large table is available at https://osf.io/kybva/.

**Supplementary Table 6. Encoded proteins of the seven main bacterial genera.** Values correspond to normalized expression for each condition (TPM+1/total number of TPM in the genus) to correct for different amount of the genus in different samples (reflected by different total TPM for the genus in different samples). Expression ratio (ER) correspond to the average of normalized expression of 9 samples at T60/72 divided by average of normalized expression of 9 samples at T0). NT: not tested if genes have less than three values for a condition in **Supplementary Table 5**. This large table is available at https://osf.io/kybva/.

**Supplementary Table 7. Encoded functions of the seven main bacterial genera.** Values correspond to normalized expression for each condition (sum of expressions for genes coding for a same function), Expression ratio (ER) correspond to the average of normalized expression of 9 samples at T60/72 divided by average of normalized expression of 9 samples at T0). NT: not tested if functions have less than three values for a condition in **Supplementary Table 5**.

**Supplementary Table 8. Transcriptomic changes of functions involved in central and energy metabolism for the different bacterial genera between T0 and the onset of oyster mortality**. Values correspond to log2 (ER). NS: not significant. Based on quantitative data presented in **Supplementary Table 7**.

**Supplementary Table 9. Differentially expressed functions in the seven bacterial genera from selected categories potentially important for successful colonization (adhesion, cell defense, metal homeostasis, redox homeostasis and oxidative stress response, stress response, and virulence/ fitness).** Based on quantitative data presented in **Supplementary Table 7.**

## Notes

### Competing Interest Statement

The authors have declared no competing interest.

### Summary of Updates

This revised version presents new analyzes on metatrascriptomics data

